# Low-latency extracellular spike assignment for high-density electrodes at single-neuron resolution

**DOI:** 10.1101/2023.09.14.557854

**Authors:** Chongxi Lai, Dohoung Kim, Brian Lustig, Shinsuke Tanaka, Brian Barbarits, Lakshmi Narayan, Jennifer Colonell, Ole Paulsen, Albert K. Lee, Timothy D. Harris

**Affiliations:** Janelia Research Campus, Howard Hughes Medical Institute, Ashburn, VA, USA; Department of Convergence IT Engineering, Pohang University of Science and Technology, Pohang, Republic of Korea; Department of Physiology, Development and Neuroscience, University of Cambridge, Cambridge, UK; Howard Hughes Medical Institute and Department of Neurology, Beth Israel Deaconess Medical Center, Boston, MA, USA; Department of Biomedical Engineering, Johns Hopkins University, Baltimore, MD, USA

## Abstract

Real-time neural signal processing is essential for brain-machine interfaces and closed-loop neuronal perturbations. However, most existing applications sacrifice cell-specific identity and temporal spiking information for speed. We developed a hybrid hardware-software system that utilizes a Field Programmable Gate Array (FPGA) chip to acquire and process data in parallel, enabling individual spikes from many simultaneously recorded neurons to be assigned single-neuron identities with 1-millisecond latency. The FPGA assigns labels, validated with ground-truth data, by comparing multichannel spike waveforms from tetrode or silicon probe recordings to a spike-sorted model generated offline in software. This platform allowed us to rapidly inactivate a region in vivo based on spikes from an upstream neuron before these spikes could excite the downstream region. Furthermore, we could decode animal location within 3 ms using data from a population of individual hippocampal neurons. These results demonstrate our system’s suitability for a broad spectrum of research and clinical applications.

## Introduction

Real-time neural signal processing is pivotal for brain-machine interfaces (BMI)^1,2^, neurofeedback therapies^3^, and closed-loop neural perturbations^4,5^. These applications rely on rapidly recognizing patterns in neural signals to trigger predefined feedback—either controlling external devices or modulating brain activity. Existing systems often compromise signal granularity for real-time processing speed: some leverage low-resolution local field potentials (LFPs)^6–11^, others over-merge^12–19^ or over-split^20–23^ spikes from individual neurons. These approaches neglect spike sorting^24–26^, a technique for identifying single neurons based on spike waveform features, often from multi-electrode arrays. Spike sorting is widely used in offline neural data analysis and provides invaluable information about coding properties (e.g., tuning curves), temporal firing patterns (e.g., bursting versus non-bursting) and cell type identities (e.g., excitatory versus inhibitory) of many individual neurons. Integrating this rich single-neuron related information, in real-time, into BMI and closed-loop neural feedback applications remains largely unexplored territory. Therefore, a low-latency neural signal processor (NSP) compatible with high-density electrodes that assigns single spikes to single neurons could impact a wide range of real-time applications.

Developing such an NSP that excels at both spike sorting accuracy and low-latency spike assignment presents a formidable challenge. Software-based NSPs lack the ability to ensure deterministic low-latency output. On the other hand, hardware solutions like field-programmable gate arrays (FPGAs), a programmable chip that can compile algorithms into digital circuits, can achieve low latency by performing data acquisition and parallel computing simultaneously. However, FPGA programming or chip design is a highly involved process requiring specialized skills. So far, on-chip spike sorting systems have only used single-channel waveform features^27–32^ or been implemented with a simplified multichannel waveform clustering algorithm^33,34^. Evidence underscores the need for sophisticated algorithms^35–37^ or even manual curation^38^ to separate multichannel spike waveform features from individual neurons and from noise. The only existing single unit, single spike, hardware-based NSPs that use multi-channel waveform features^33,34^ have yet to prove their ability to de-noise data and accept user-defined spike sorted models—software features often essential for in vivo data analysis. Lacking flexible software, these pure on-chip solutions have not been deployed in laboratory research applications.

Here, we introduce a hybrid hardware-software system that harnesses the strengths of both the speed of hardware and the flexibility of software. Utilizing a previously proposed two-stage method^17^, we first establish a spike-sorted model in software, then transfer the model parameters to our FPGA-based NSP for real-time spike detection and assignment. The design delivers flexibility speed, assigning single-neuron identities to all detected single-spikes with 1-millisecond latency across 160 channels^39^, and has recently been deployed in online experiments^40^. Unlike prior reports^27–34^, our platform accommodates multichannel spike waveforms and is compatible with tetrodes and silicon probes, while enabling modern clustering algorithms and manual curation for crafting accurate spike-sorted models. To abstract the hardware complexities, we provide a backend Application Programming Interface (API) and frontend modular Graphical User Interface (GUI) components for intuitive data visualization, data management, model curation, and FPGA-NSP status control. We also provide an Open Ephys plugin such that users can use Open Ephys acquisition software^41^ with our system.

## Results

### System design and the FPGA-based NSP

Our system leverages a commercially available Xilinx KC705 FPGA board paired with a 48-core PC workstation, running our Spiketag^42^ software package (Fig. 1a, b). The FPGA board is connected to the workstation via a Peripheral Component Interconnect Express (PCIe) slot. The FPGA-based NSP processes raw data from 160 electrodes, amplified and digitized at 25 kHz by five 32-channel INTAN RHD2132 acquisition chips. As the data flows into the FPGA’s algorithmic circuits (Fig. 1b), real-time neural signal processing starts. The first step in the processing pipeline involves noise filtering and artifact removal through reference subtraction (Fig. 1b). This effectively filters out both low-frequency (<500 Hz, i.e., LFP) and high-frequency (>3000 Hz, i.e., noise) components and eliminates motion artifacts commonly present across channels (Fig. 1c, d).

**Fig. 1.**
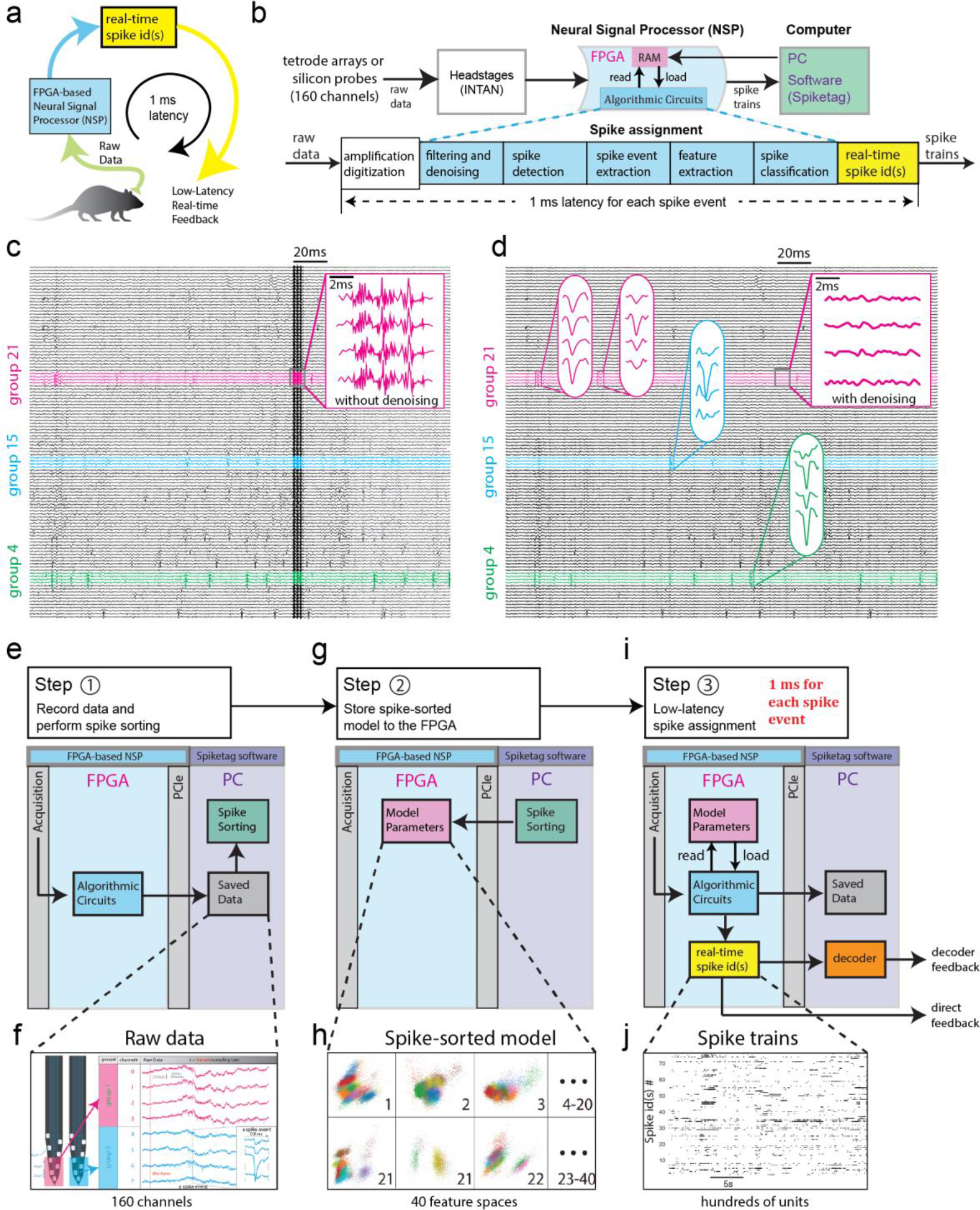
Hybrid hardware-software system for low-latency assignment of spikes to individual neurons. Our system outputs real-time spike trains by assigning each spike a spike-id based on a spike-sorted model. **a**, The FPGA-based NSP takes the raw data and outputs the real-time spike ids, that can be used to trigger the low-latency feedback. The spike assignment loop takes less than 1 ms for each spike event. **b**, An expanded view of (a). The real-time signal processing of our FPGA-based NSP relies on our Spiketag software. Spiketag provides a set of Python APIs to acquire and parse preprocessed data from the FPGA-based NSP and download a set of parameters into the FPGA’s memory (RAM) to guide hardware-implemented algorithms such as filtering, denoising, spike detection, multi-channel spike extraction, feature extraction by a linear transformation (e.g., PCA), and spike classification (collectively named “Algorithmic Circuits” as all of these algorithms are implemented in digital circuits). These FPGA-based NSP algorithmic modules were pipelined for spike assignment. **c**, Filtered but not denoised multi-channel signals (left: the motion noise is similar across all channels as shown inside the black rectangle), and **d**, Filtered and denoised multi-channel signals of the same recording segment (right). Data was partitioned into groups of four electrodes each for independent processing. Close-ups of four examples of four-channel spike events from three different electrode groups (in green, blue, and pink) are shown. **e**, The first step when using our system is to record some data and perform rapid spike sorting (clustering), which results in a spike-sorted model. **f**, Examples of 8 channels of raw data, representing 2 electrode groups. **g**, The second step is to store the spike-sorted model to the FPGA. **h**, Example of stored model in the FPGA (before vector quantization). **i**, Using this model in the FPGA, the NSP performs low-latency data processing and outputs a spike-id for each spike. **j**, Example of real-time sorted spike trains output during recording.

After filtering and noise removal, the 160 electrodes are separated into 40 non-overlapping electrode groups, each containing four adjacent electrodes, for spike detection and extraction (Fig. 1b, see three electrode group examples highlighted in Fig. 1c, d). Spike detection is initiated by a threshold-crossing event. A 0.76 ms-wide snippet of neural activity is captured. This snippet is combined with coincident spikes from other channels in the same electrode group to form a multichannel spike event (Fig. 1d). Each detected spike event then undergoes transformation and embedding into the corresponding feature space computed from pre-recorded raw data. Using k-nearest neighbor (kNN) classification, each spike is assigned a unique spike-id based on the label of its nearest cluster (of the spike-sorted model that was determined using pre-recorded data) in that feature space.

kNN classification was chosen as it is one of the most accurate algorithms for on-chip spike assignment^27^, given that waveform clusters can be non-Gaussian^43^ and may not be linearly separable. However, standard kNN would require a large memory for FPGA implementation. Here, vector quantization was used to compress the thousands of points in each single cluster into a few tens or hundreds of points (reference vectors), saving FPGA memory usage and reducing spike classification time by several orders of magnitude, making this version of kNN FPGA-implementable with similar performance to standard kNN.

Spiketag allows users to transform sample recordings into spike-sorted models (Fig. 1e, f), parameters of which are then downloaded into the FPGA for real-time spike assignment (Fig. 1g, h). The model parameters include the channel grouping, the reference channels, the spike detection thresholds, the feature transformation, and vector-quantized feature clusters of each group. Once downloaded, the NSP switches to ‘inference mode,’ automatically generating spike trains assigned to single neurons (clusters) with a guaranteed low latency of 1 ms for each spike, irrespective of the number of neurons (Fig. 1i, j)

As described in the introduction, by allowing the hardware (NSP) and software (Spiketag) to communicate seamlessly, our system simultaneously achieves low latency and accurate spike assignment according to a user-defined spike-sorted model. Users can flexibly decide the level of the required accuracy of single-unit assignment for their experiment, ranging from under-clustered (i.e., sorting into multi-units), to accurately clustered (into single units), to over-clustered (i.e., over-split or “clusterless”) models. We optimized our software to rapidly generate such coarse to accurate spike-sorted models through fast 3D visualization and interactions.

### Spiketag enables interactive spike sorting, NSP parameter adjustment, and visualization of NSP-processed data

The ability to rapidly perform visual inspection and manual curation of the spike-sorted model is critical for real-time experiments. Over time, electrode drift can compromise the model’s accuracy. Some electrodes might become excessively noisy. Such scenarios necessitate rapid parameter adjustments. To allow rapid adjustment of NSP parameters and visualization of its effect, Spiketag software was designed with two key features. First, we separated the software into a frontend and backend. The frontend provides several graphical widgets to interact with data visually while the backend performs data analysis and automatic clustering for groups in parallel using different CPU cores (Fig. 2). Second, enabled by Vispy^44^, a high-performance interactive 2D/3D data visualization library, we developed a series of interactive visualization modules in Spiketag which respond to user input instantaneously. Users can thus navigate through all electrode groups via a Probe View using a mouse and keyboard, and select one electrode group for manual curation. While a user curates one group, the backend applies the clustering algorithm to other groups. The visually-based data interaction also allows users to perform manual curation rapidly. For example, users can use the mouse and keyboard to split or merge clusters in the Feature View or Spike View and see instant updates in all of the other views (Fig. 2e). Meanwhile, Spiketag’s backend automatically updates the model parameters based on user input that the user can subsequently download to the NSP.

**Fig. 2.**
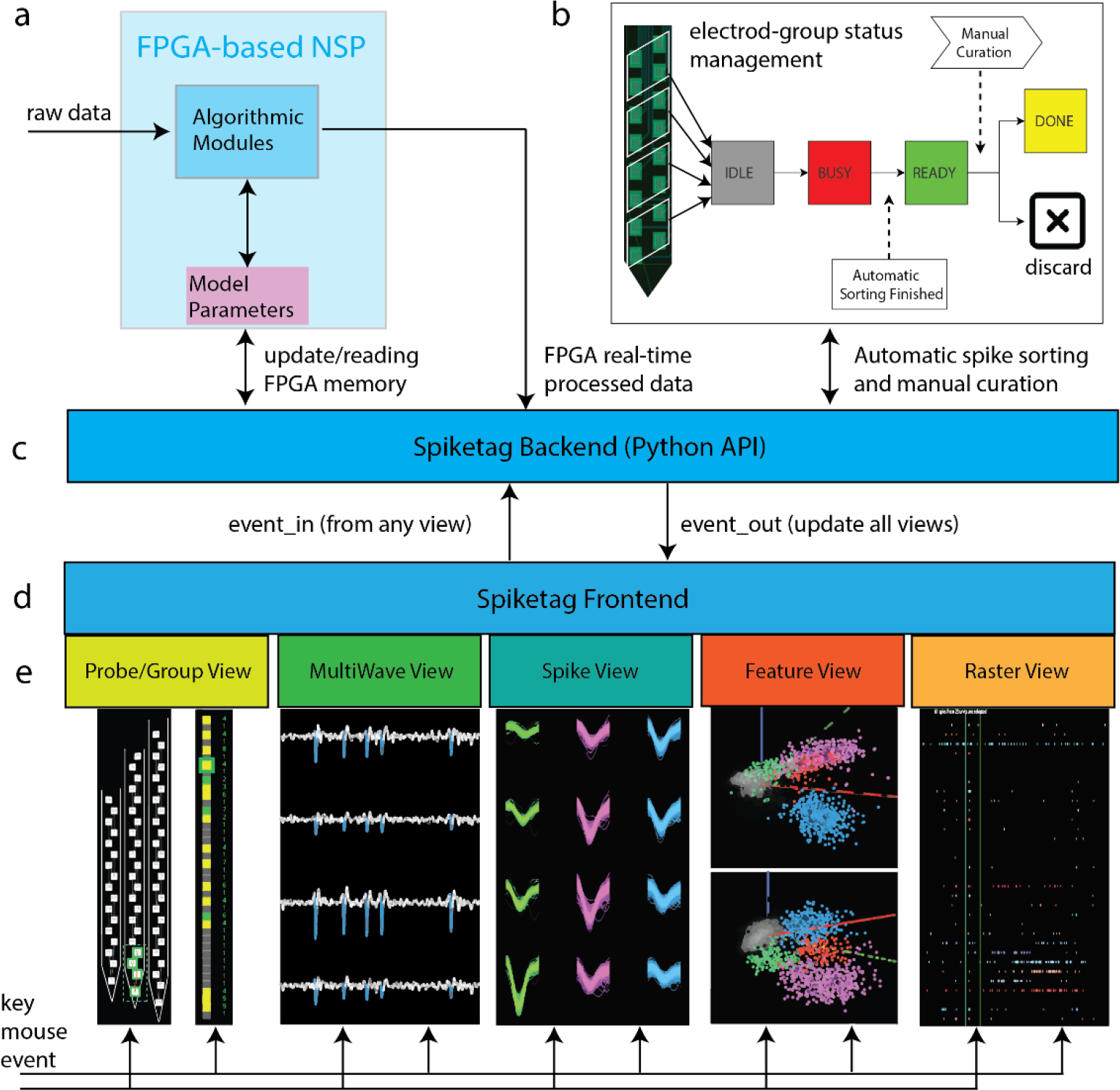
The Spiketag software package allows users to update parameters in the NSP based on the spike-sorted model created in the PC. The model is generated in the PC from automatic spike sorting followed by manual curation. **a**, The Spiketag backend can read and write the model parameters inside the FPGA to steer the online processing. It also receives real-time FPGA processed data, used for real-time visualization by the frontend (d). **b**, The operating status of each electrode group is visually represented by their colors in the Probe/Group View (e), so that the user knows which electrode group contains spikes and which electrode groups have completed automatic clustering and are ready to be visually inspected. The automatic clustering is scheduled in idle CPUs by the backend. **c**, The backend was developed with a set of low-level APIs to handle FPGA communications, read and save FPGA real-time processed data, schedule automatic sorting, and communicate with the Spiketag frontend. **d, e**, The Spiketag frontend was developed with several Vispy-based interactable views to present different types of data in both online and offline mode. When interacting with any of these views, the keyboard or mouse input is converted into an event and is relayed to the backend (event_in), and the backend in turn process it and sends commands (event_out) to other relevant views to update them.

Spiketag places users in the loop by providing visual feedback and allows users to interactively adjust parameters to produce verifiable intermediate partial results. These parameters include the spike detecting thresholds of any electrode, reference electrodes for canceling noise, features transformations, and clusters, which in turn progressively give rise to the end result of spike trains. For example, to prevent false positive spikes detected from a noisy electrode, all that is required is for the user to raise the spike threshold for that channel to a very high value; to exclude an unstable or unwanted cluster in a feature space, the user need only mark that entire cluster as noisy (i.e., cluster 0). By adjusting these parameters, the NSP is progressively configured to generate the desired output. The above steps are generally done in offline mode to create the spike-sorted model. Spiketag also supports an online mode in which real-time data streams into the views and old data gradually fades away. The online mode allows users to closely monitor the processing of data by the NSP, including spike assignment, during a perturbation or BMI experiment.

Spiketag also provides a set of Python APIs that allows users to build upon the real-time processed data described above. Spiketag modules can be imported into the user’s Python code. For instance, it is often a critical part of an experiment for users to evaluate the receptive fields of the sorted neurons (which can be done in a Jupyter notebook), or connect the real-time spike trains to a decoder for BMI control or perturbations (e.g., see population decoding section below). In addition to spike train data, users can analyze all processing steps of the NSP, including filtering, noise removal, spike detection, feature extraction, clustering, and vector quantization.

### Verification of latency and spike sorting accuracy

In order to test the accuracy and latency of our NSP’s real-time spike assignment, it is necessary to use ground truth data and test the latency from raw analog input to the final spike-id output. We built a custom neural signal generator (NSG) to replay up to 192 channels of simulated or pre-recorded extracellular traces as analog signals between tens to hundreds of ±μV at 25 kHz, as in real neural recordings (full output range ±5 mV). 160 of the 192 NSG analog outputs were fed into the five INTAN chips of our system as in a live experiment. A TTL signal output by the NSG (on channel 161) indicating the ground-truth spike times, and a TTL triggered by the FPGA upon spike-id assignment, were both fed into a logic analyzer running at 50 MHz to compute the latency. The measured latency of the device under test (DUT) thus includes the delay introduced by the INTAN chips and our NSP chip (Fig. 3a-d). To test the latency, we synthesized 161-channel data containing four artificial ground-truth neurons firing at a fixed interval with realistic spike waveforms across several channels (Fig. 3e). These neurons were accurately detected and assigned ids without error. One neuron that had the largest spike amplitude on channel 158 was selected as the trigger unit whose spikes were used to trigger an NSP output TTL (i.e., spike assignment-triggered TTL). The latency was calculated by comparing the ground-truth TTL and the spike assignment-triggered TTL (Fig. 3e). The result shows that when an extracellular spike waveform comes to completion, its identity will be accurately assigned in 0.973±0.012 ms (Fig. 3e, f). This latency includes the delay from signal amplification, filtering, spike detection, and all other preprocessing steps.

**Fig. 3.**
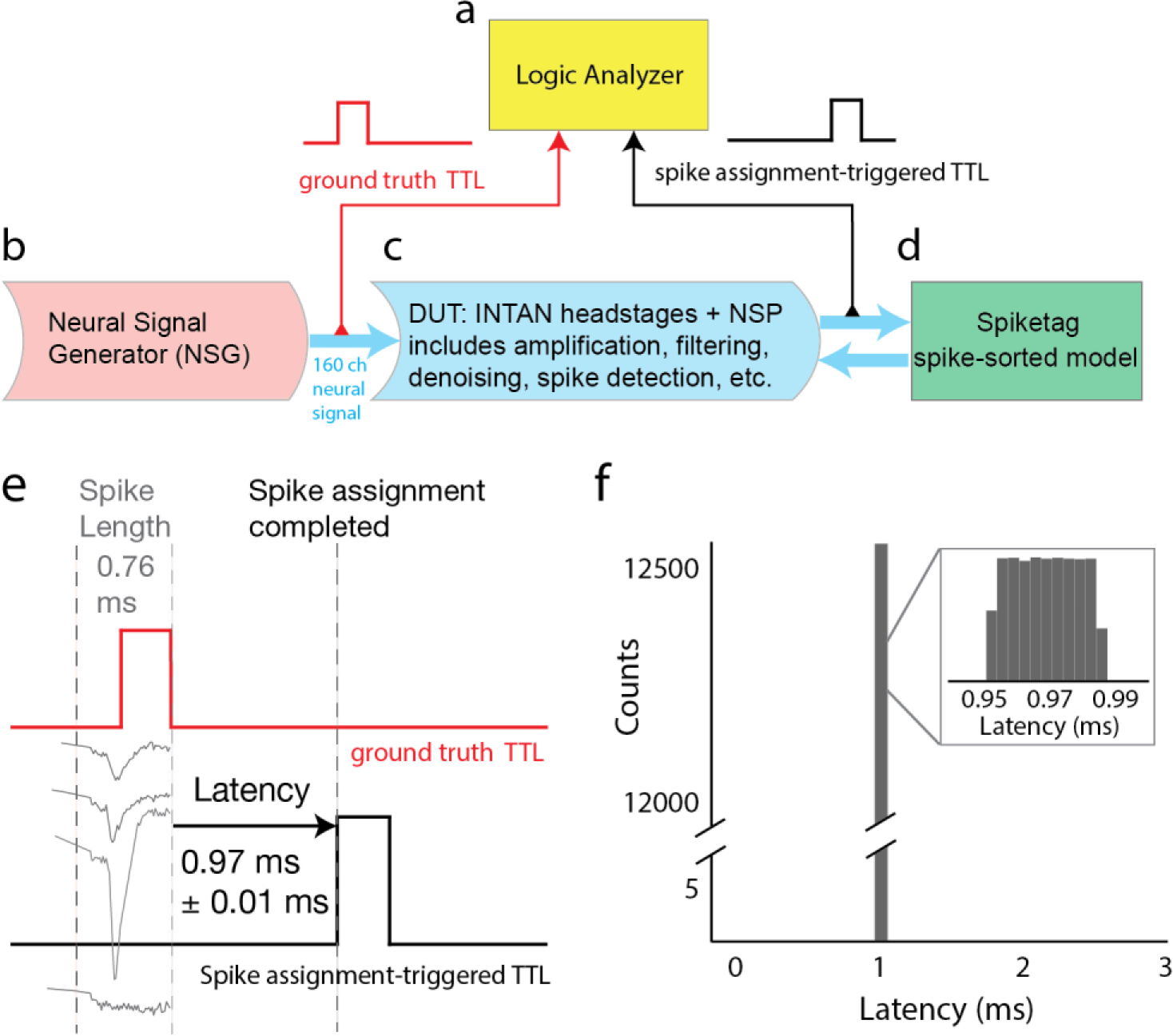
The FPGA-based NSP classifies the source of each spike within 1 ms. Latency test for single-unit spike assignment. **a**, A 50 MHz logic analyzer receives both ground-truth TTL and the FPGA spike assignment-triggered TTL. **b-d**, The NSG outputs a ground-truth TTL signal using one extra channel (a 161st channel), indicating the timing of the ground-truth spikes, to the logic analyzer. The device under test (DUT) includes the INTAN headstages and the FPGA NSP. The NSP outputs a TTL triggered by spike assignment to the same logic analyzer. **e**, An example of the ground-truth TTL (upper, red) and the spike-triggered TTL (lower, black). The 0-1 transition of the ground-truth TTL indicates the peak sample of the largest spike waveform across all four channels, and the 1-0 transition indicates the end of the spike waveform. The spike-triggered TTL (black) indicates this neuron was identified to be the trigger unit. Its transition from 0-1 reveals a 0.973±0.012 ms latency from spike completion (in the raw analog signals) to its correct assignment by the NSP. Importantly, this latency includes the delay from signal amplification, filtering, denoising, spike detection, and all other preprocessing steps to the final spike-id assignment. **f**, The histogram of the latency as illustrated in (e). The latencies remained stable during a 33.8 minute test and spanned 0.952 ms to 0.998 ms with zero outliers found outside this range.

To test the accuracy of our spike sorting method, we applied two previously published ground-truth datasets^45,46^ to the same test system described above. Both datasets contain a ground-truth neuron whose spikes are known. For each dataset, we generated a spike-sorted model, downloaded it to the FPGA-NSP to perform real-time spike assignment, compared the corresponding single unit’s spike train to the ground-truth spike train, and measured accuracy in terms of hit rate and precision. The first dataset was collected using a tool called a Patch-Tritrode, which allows the placement of three extracellular electrodes a few tens of μm away from an intracellular patch pipette electrode^46^. A sparse and bursty hippocampal CA1 pyramidal neuron was simultaneously recorded intracellularly (ground-truth) and extracellularly. Both the hit rate and precision of our low-latency assigned spike-ids were as high as other state-of-the-art offline sorting packages^36,37^, even for this sparse and bursty neuron. The second dataset involved simultaneous juxtacellular and extracellular recording of a neuron by a patch pipette and custom high-density silicon probe in the primary motor cortex^45^. Because the silicon probe has 32 electrodes, four electrode groups near the ground-truth neuron were created and selected, each containing four electrodes covering an area of 60 μm by 50 μm on the probe. Our system correctly assigned the ground-truth neuron’s spikes with high accuracy (Table 1). Therefore, our NSP can deliver single-unit spike assignment accuracy comparable to offline spike sorting methods for both tetrode and silicon probe data.

**Table 1.** Accuracy of single-unit inference on ground-truth data. Two ground-truth datasets (from a tritrode and silicon probe) were used to test accuracy in terms of hit rate and precision. We compared our accuracy on the ground-truth dataset 1 with Kilosrt1 and JRClust. Four electrode groups were created on the silicon probe for ground-truth data 2. Since our system processes waveforms separately from each electrode group whenever a spike is detected, the hit rate and precision was calculated for the same neuron for each of these groups.

| Ground-truth dataset 1<br>From Tritrode |  | Hit rate | Precision |
| --- | --- | --- | --- |
|  | This system | 93% | 87% |
|  | Kilosort 1 | 88% | 92% |
|  | JRClust | 96% | 83% |
| Ground-truth dataset 2<br>From silicon probe |  | Hit rate | Precision |
|  | This system<br>Electrode group #0 | 99.7% | 97.1% |
|  | This system<br>Electrode group #1 | 98.2% | 95.3% |
|  | This system<br>Electrode group #2 | 96.4% | 98.8% |
|  | This system<br>Electrode group #3 | 97.1% | 93.2% |

### Fast closed-loop perturbation triggered at single-unit single-spike resolution

An advantage of our low-latency, single-unit, single-spike assignment is the potential to perturb a downstream area before specific spikes from an upstream area arrive. Here we investigated this possibility by recording from the lateral geniculate nucleus (LGN) and optogenetically suppressing (via activation of local GABAergic neurons) the primary visual cortex (V1) in a transgenic VGAT-ChR2-eYFP mouse. One 64-channel silicon probe was implanted in LGN and another in V1 (Fig. 4a). An optical fiber was attached to the V1 probe to perform optogenetic stimulation (Fig. 4a). The awake, head-fixed mouse was placed in front of a visual display during recording. After spike sorting a 10 min recording, one of the LGN neurons was selected as the source neuron (Fig. 4c,d). Then the spike-sorted model was downloaded into the FPGA-NSP to guide real-time spike assignment. The FPGA-NSP output a TTL pulse whenever the source neuron fired a spike to drive the laser. Half of the trials were randomly selected for the closed-loop stimulation. Each trial started with a visual stimulus. During closed-loop trials, each spike from the LGN source neuron triggered the laser for 20 ms to suppress V1. Some laser durations were elongated due to additional source neuron firing within this 20 ms period (see distribution in Fig. 4g).

**Fig. 4.**
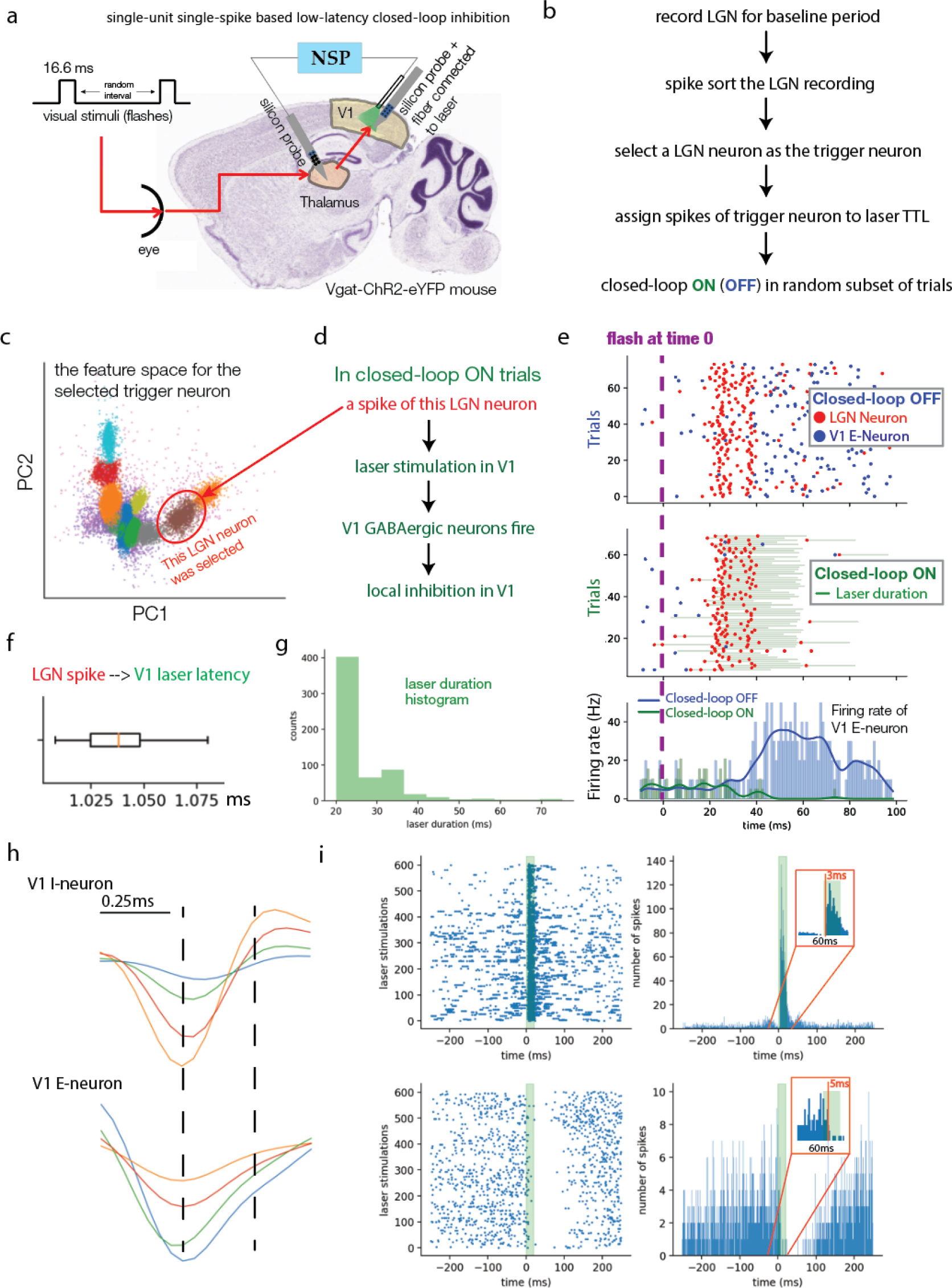
Rapid suppression of a downstream region based on upstream activity. LGN single-unit spike-based closed-loop perturbation of V1: **a**, Schematic of the experiment. The LGN and V1 were recorded simultaneously while LGN activity was processed online. **b**, Experiment steps. After recording a baseline period (10 min) in LGN and creating the spike-sorted model in Spiketag, a source neuron was selected manually so that when it fires a spike the NSP outputs a TTL pulse which drives a laser for photoinhibition in a random subset of trials. **c**, The feature space of the group from which the LGN source neuron (circled) was selected. **d**, Laser stimulation causes V1 GABAergic neurons to fire within 3 ms, leading to local inhibition in V1 (e). **e**, An example of the LGN source neuron (spikes indicated by red dots) and a V1 excitatory neuron (V1 E-neuron, spikes indicated by blue dots) firing during non-closed-loop (upper) and closed-loop trials (middle). This plot is aligned by trial, and each trial starts with a flashing visual stimulus at time 0. In non-closed-loop trials (upper), the LGN source neuron mainly fires (red dot) between 20 ms and 40 ms, immediately followed by an increased firing rate of a V1 E-neuron (blue). In closed-loop trials (middle), the same LGN source neuron’s spikes lead to optogenetic inhibition in V1 (20 ms pulse triggered upon each spike, total laser durations are indicated by green lines). The optogenetic inhibition blocks the vast majority of V1 E-neuron spiking and the stimulus-triggered increase in firing rate. The firing rate of the same V1 E-neuron in non-closed-loop trials and closed-loop trials are compared (lower). **f**, The LGN spike from the source neuron to V1 laser stimulation takes ∼1 ms. Latency was measured by an independent acquisition system (NI) that receives timing signals from the NSP and laser system. **g**, Histogram of laser durations. While each spike triggers 20 ms of laser stimulation, multiple source neuron spikes firing within 20 ms can stretch the laser duration. **h**, Spike waveforms, recorded from a single electrode group, were classified as V1 inhibitory neuron (I-neuron) and V1 excitatory neuron (E-neuron) according to their spike width. **i**, The peri-stimulus histogram (aligned to laser onset timing) of both the V1 I-neuron and the V1 E-neuron in (h). The LGN source neuron spike-induced laser duration (20 ms) is highlighted in green. The insets show the histogram from –30 ms to 30 ms. Aligned to the laser onset, they show that the inhibitory neuron increased its firing rate within 3 ms and the excitatory neuron decreased its firing rate within 5 ms.

In our experiment, an excitatory V1 neuron (blue dots in Fig. 4e) consistently increased its firing in response to the stimulus in the absence of low-latency optogenetic silencing (Fig. 4e Closed-loop OFF panel). In contrast, this neuron stopped firing in the vast majority of closed-loop trials in which optogenetic silencing was triggered by LGN source neuron spikes and, in particular, the stimulus-evoked increase in firing rate disappeared (Fig. 4e Closed-loop ON panel). In trials in which the LGN trigger neuron fired (red dots in Fig. 4e Closed-loop ON panel), the laser was triggered in ∼1 ms (Fig. 4f). The laser led to increased firing of GABAergic neurons in ∼3 ms (Fig. 4 h, i, consistent with a previous study^38^). Together, our effective closed-loop perturbation latency (from LGN spike to V1 inhibition) via photoinhibition was ∼4 ms. This latency (4 ms) is comparable to the monosynaptic EPSP latency from LGN to V1 measured previously (4.9 ± 0.5 ms)^47,48^, and shorter than the monosynaptic latency of other pathways^49^, thus allowing us to suppress a downstream area before spikes from an upstream area can excite it. Such spike-triggered cross-region perturbation would not be feasible without a low-latency NSP. This experimental paradigm could also be applied to perturb a downstream region based on the combinatorial spiking pattern of a population of single units in the upstream region, either to intercept information transmission between brain areas via inhibition or to introduce plasticity between neural assemblies via excitation.

### Fast, accurate real-time population decoding

We next verified the accuracy and latency of population decoding with our system. The test dataset was hippocampal CA1 activity recorded using silicon probes from a freely moving rat that randomly foraged in a 160 cm by 160 cm arena. Individual hippocampal neurons display location-specific (place field) activity^50^. The activity of a population of such neurons can be used to decode the location of the animal, which is usually performed offline. Similar to the previous test configuration, 160 channels of raw data were replayed as analog signals by the NSG, and neuronal sources of spikes were assigned in real-time by the NSP then transmitted to the computer via PCIe for real-time location decoding. A Python script was used to receive NSP spike-id output and decode animal position in real-time. Reference TTL pulses (every 40 ms) were encoded as spike times of an artificial neuron. The Python script started decoding when it received the spikes indicating the reference TTL pulse. Once the decoding was finished, the Python script triggered a microcontroller (Teensy) connected to the PC to produce a feedback TTL pulse. The reference and feedback TTL pulses were fed into the logic analyzer to measure the full processing latency (Fig. 5a). While the previous section tested the latency from raw spike data to spike-id assignment, here the latency also included transmitting the spike-id packets from NSP to PC, the PC receiving the spike-id packets, PC population decoding, and the PC generating the feedback TTL signal.

**Fig. 5.**
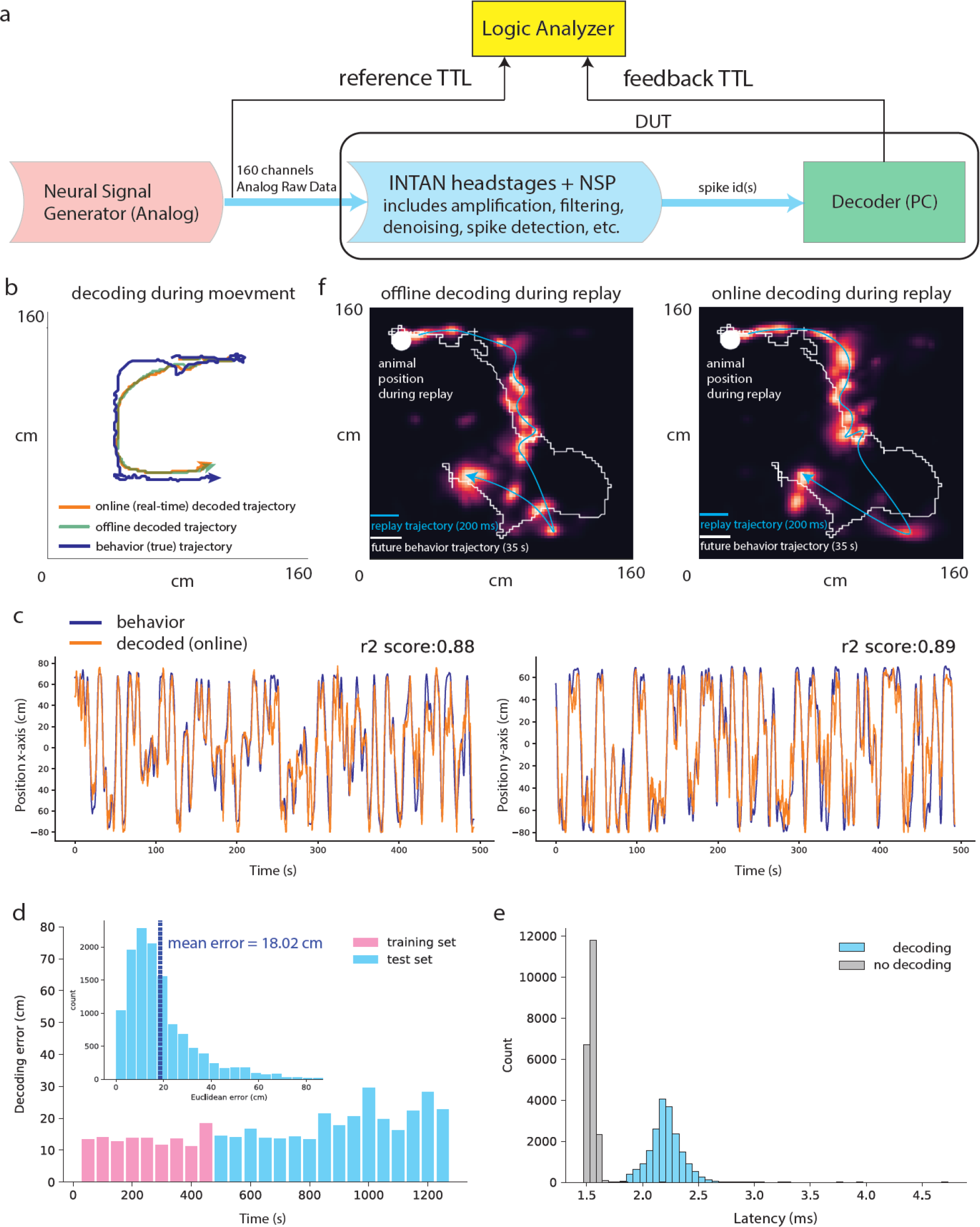
Rapid and accurate online population decoding of animal trajectory. Tests were performed using the hippocampal activity of a population of CA1 place cells simultaneously recorded from a freely moving rat running in a square-shaped arena. **a**, Configuration to test the latency of population decoding by comparing the timing of reference and feedback TTLs. The DUT here includes the decoder running on a PC. **b**, An example showing the past 30 s of actual (blue), online decoded (orange), and offline decoded (green) locations of the animal. Arrows indicate the movement direction. **c**, The overlayed traces of the online decoded trajectories (orange) and the actual trajectories (blue) of the test (i.e., cross-validation) data (left: x-axis, right: y-axis). Decoding results during periods that the rat did not move were excluded. **d**, The decoding error distribution of both the training (magenta) and test (cyan) sets. Each bar represents 50 s. Inset: The distribution of decoding error (cyan) over the full test set. **e**, Histogram of the full processing latency (raw data to amplifiers, A/D conversion, NSP acquisition and single-unit assignment, using spike-ids to decode animal positions, then final output of the feedback TTL pulse). This latency was tested with and without decoding (gray vs cyan). **f**, A representative replay event decoded offline (left) and online (right) overlayed across time (200 ms) with a 25 ms decoding window advancing every 5 ms. The replay event occurred when the rat was still (animal position: the white dot). The replay trajectory (cyan line) was interpolated using positions of maximum posterior probabilities across 200 ms (overlayed on a heat map of decoded posterior) and predicts 35 s of immediate future movement trajectory (white line). The slight, expected difference between online and offline decoded heat maps is due to the difference in spike-sorted models. The offline model uses all the data, while the online model was determined from training data only.

The place fields of 76 neurons were constructed offline from a validation dataset (first 30% of the full recording while the animal’s speed was >10 cm/s) and then used with a memoryless Bayesian decoding algorithm^51^ to convert real-time population spike trains from the test dataset (the last 70% of the data) into specific locations in the 2D arena. A 280 ms decoding window was used to generate each decoded location and advanced in 40 ms steps. The sequence of decoded positions was concatenated into a trajectory and smoothed. The online decoded and smoothed trajectory closely tracked the animal’s actual trajectory in the validation set (Fig. 5b-d). The test results showed an average 18.0±12.4 cm (Fig. 5d inset) decoding error. The mean decoding error remained stable across the validation set, shown by separating the validation set into 50 s bins (Fig. 5d). The decoded results were generated with latency 2.2±0.15 ms after the reference TTL pulses (Fig. 5e). The decoding computation itself contributed modestly to the latency. If decoding was skipped, the latency to produce the feedback TTL was 1.55±0.04 ms (Fig. 5e).

This population decoding feedback latency is smaller than the steps usually used to decode fast trajectories during hippocampal replay^35^ (3-5 ms), therefore our system can be used for feedback based on rapidly changing replay content. For instance, analyzing the real-time decoding of hippocampal replay content using a 25 ms decoding window advancing every 5 ms, we observed many replay trajectories during periods when the rat was stationary that predicted the animal’s future movement trajectories (example shown in Fig. 5f), as reported previously^52^.

Overall, our system can decode 2D replay trajectories in real-time as accurately as offline decoding, achieving low-latency performance while providing single-unit specificity. Moreover, users can flexibly create an over-split spike-sorted model, akin to clusterless decoding^20–23^, in real-time applications where single neuron information is not required. This can reduce the time spent spike sorting, which can be critical for fast model deployment. Recently, we used the real-time population decoding capabilities of this system to demonstrate that an animal can volitionally control its hippocampal place-related activity using a BMI^40^.

## Discussion

We report a high-performance tool capable of assigning single-spike activity from individual neurons with a latency of 1 ms during multi-electrode data acquisition. Our system supports up to 160 channels, is versatile, and compatible with any recording system employing affordable, commercially available INTAN digital headstages. It can be scaled for higher electrode channel counts by integrating multiple systems. Using ground-truth data sets, in vivo closed-loop optogenetic experiments, and in vivo population recordings, we demonstrated the system’s speed, precision, and applicability to real-world scenarios, from closed-loop perturbations to population decoding.

Our tool has transformative potential in the realm of brain-machine interfaces (BMI) and closed-loop experiments. Traditional BMIs typically focus on population rate coding, capturing only the firing rates of multi-unit activity. Our tool takes this a step further by enabling the design of BMI decoders based on temporal coding, allowing for real-time detection of complex spike patterns and neuronal bursting events. Moreover, our platform provides cellular specificity, essential in experiments where gathering cell-type information is critical for dissecting and perturbing neural circuits. Combined with optogenetic tagging and spike waveform classification^53^, our tool allows for real-time, cell type-specific, and spike-based interventions, setting it apart from other existing methods for dissecting neural circuits and probing the function of putative neural codes. For instance, with a 1ms latency, our platform can investigate the causal roles of hypothetical neural codes via the suppression of downstream effects of specific activity patterns. Additionally, this system can be used for exploring neuronal plasticity rules with high temporal and cell-type specificity across multiple neurons. In conclusion, our system provided unprecedented resolution and speed for both BMI and closed-loop neural perturbation experiments, opening up possibilities for a variety of new neuroscience applications.

## Acknowledgments

We thank Nakul Verma for valuable discussions regarding algorithms for compressing clusters, Aarón Cuevas López for valuable discussions regarding INTAN SPI firmware, Jim Chen for valuable discussions on Xilinx FPGA programming, and Yixin Chi for technical assistance with Spiketag software programming. This work was supported by the Howard Hughes Medical Institute.

## Notes

### Competing Interest Statement

The authors have declared no competing interest.

